# *A Day in the Life of Arabidopsis:* 24-Hour Time-lapse Single-nucleus Transcriptomics Reveal Cell-type specific Circadian Rhythms

**DOI:** 10.1101/2023.12.09.570919

**Authors:** Yuwei Qin, Zhijian Liu, Shiqi Gao, Yanping Long, Xinlong Zhu, Bin Liu, Ya Gao, Qiguang Xie, Maria A. Nohales, Xiaodong Xu, Jixian Zhai

## Abstract

Functional circadian clock is fundamental to the adaptation and survival of organisms. In land plants, the comprehensive profiling of circadian gene expression at the single-cell level is largely unknown partly due to the challenges in obtaining precisely-timed single cells from plant cells embedded within cell walls. To bridge this gap, we employed single-nucleus RNA sequencing (snRNA-seq) on twelve seedling samples collected at 2-hour intervals over a 24-hour day in Arabidopsis, yielding a total of over 130,000 nuclei. From this data, we found that three cell clusters in the shoot share a coherent rhythm, while more than 3,000 genes display cell-type specific rhythmic expression. Only 19 genes are oscillating in more than ten different cell types, and the majority of them are well-documented core oscillators, suggesting the snRNA-seq circadian data could be used to identify key circadian regulators in a broad range of plant systems. Our data provides the first comprehensive resource for plant circadian rhythms at the single-cell level (hosted at https://zhailab.bio.sustech.edu.cn/sc_circadian).

## Introduction

The circadian clock is a time-keeping mechanism that coordinates 24-hour rhythms synchronized with environmental cycles, which is essential for the temporal organization of physiology in organisms^1–3^. The core oscillator consists of the transcriptional-translational feedback loops (TTFLs), which regulates gene expression from dawn till late night, driving metabolism and growth in synchronization with the day-night cyclic cues^4–7^. Within plant systems, the circadian clocks regulate distinct rhythmicity and functions among different tissues and cell types^8–13^.

In plants, studies on circadian rhythms have extensively utilized techniques such as bioluminescence tracking of *promoter:LUC* and real-time quantitative PCR (qRT-PCR) to monitor the promoter dynamic activity of the reporter genes or the transcripts rhythmic enrichment in whole seedlings and various tissues^14–19^. Despite these findings, these studies obscured the differences in circadian rhythms at the individual cell level. Previous studies using long-term imaging approaches have provided glimpses into this complexity^18,20–26^. However, these methods, while informative, have lacked the throughput to comprehensively dissect the operation of the clock system at the single-cell level.

The advent of single-cell RNA sequencing (scRNA-seq) has revolutionized our understanding of cellular heterogeneity, providing a lens to view gene expression dynamics at an unprecedented resolution^27–29^. In recent years, scRNA-seq has been used to delineate circadian gene expression in animals, providing insight into the temporal regulation within individual cells with samples collected in four-hour intervals^30,31^ or at two time points^32,33^. A recent study in plant also used scRNA-seq to examine protoplasts of Arabidopsis shoots and roots collected at end of day or end of night^34^. However, plant cells, encased by rigid cell walls, pose unique challenges for short interval time-series analysis, because the protoplasting process required for plant single cell isolation can take several hours and therefore unideal for capturing the time-series transient transcript changes of circadian regulation. To overcome issues induced by protoplasting, a recent study by Torii et al. manually extracted cellular content from 216 single cells collected at different time points using microcapillary and constructed scRNA-seq library for each cell, and their results shed light on the pivotal role of the circadian clock in leaf cell differentiation processes^35^. However, a comprehensive atlas of single-cell circadian gene expression for a whole Arabidopsis seedling remains to be developed.

In this study, we have overcome these limitations by employing single-nucleus RNA sequencing (snRNA-seq), a technique less impeded by the physical constraints of plant cell walls. By applying snRNA-seq to Arabidopsis seedlings, we have mapped circadian gene expression in different cell types with high temporal resolution at two-hour intervals. Our results reveal significant oscillator similarities among three shoot cell clusters and identify approximately 3,000 genes that oscillate within a specific cell cluster. Furthermore, we have detected only about 19 genes oscillating in more than 10 cell types, the majority of them being well-known clock genes, highlighting the potential of using snRNA-seq to uncover new clock components in plant organisms.

Through this comprehensive dataset, we provide an invaluable resource for further investigation into plant circadian rhythms at the single-cell level, offering a new dimension of understanding how temporal regulation can influence plant biology at the most fundamental level. Our findings represent a considerable step forward in plant circadian biology, promising to enrich our understanding of plant physiology and potentially guiding agricultural practices to harness the innate timing mechanisms of crops for optimized growth and yield.

## Results

### Single-nucleus RNA-seq reveals circadian rhythms in Arabidopsis

To craft a detailed single-cell transcriptomic blueprint of temporal mRNA rhythmic accumulation via the circadian clock, we leveraged the 10X Genomics platform to execute snRNA-seq on whole seedlings, harvested every 2-hour across 12 circadian time points under constant light (LL; free-running conditions), with the sequencing libraries constructed in two batches (Fig. 1A). After performing quality control and computing the expression matrix, we acquired a dataset of 131,152 high-quality single-nucleus transcriptomes spread across twelve libraries, encompassing 24,503 genes, with an average of 847 genes and 1,058 UMI detected per nucleus (Supplementary Table 1). Our data revealed pronounced circadian rhythm patterns, with increased transcriptomic similarity between adjacent circadian time points, regardless of the batch they originated from (Fig. 1B). This confirmed that our approach effectively captured the dynamic changes in clock gene expression. To further validate our methodology, we employed a pseudo-bulking approach to aggregate gene counts in each sample. As expected, the expression patterns of 17 core clock genes^7^ (*CCA1*, *LHY*, *RVE4*, *RVE8*, *RVE6*, *LNK1*, *LNK2*, *PRR9*, *PRR7*, *PRR5*, *PRR3*, *TOC1*, *GI*, *LUX*, *ELF4*, *ELF3* and *ZTL*) showed strong rhythmic oscillations peaking at anticipated time points (Fig. 1C), such as *CCA1* and *TOC1*, peaking at subjective dawn and dusk, respectively (Fig. 1D). Collectively, these findings underscore the feasibility of our approach in analyzing circadian rhythms using single-nucleus transcriptional profiling.

**Fig. 1:**
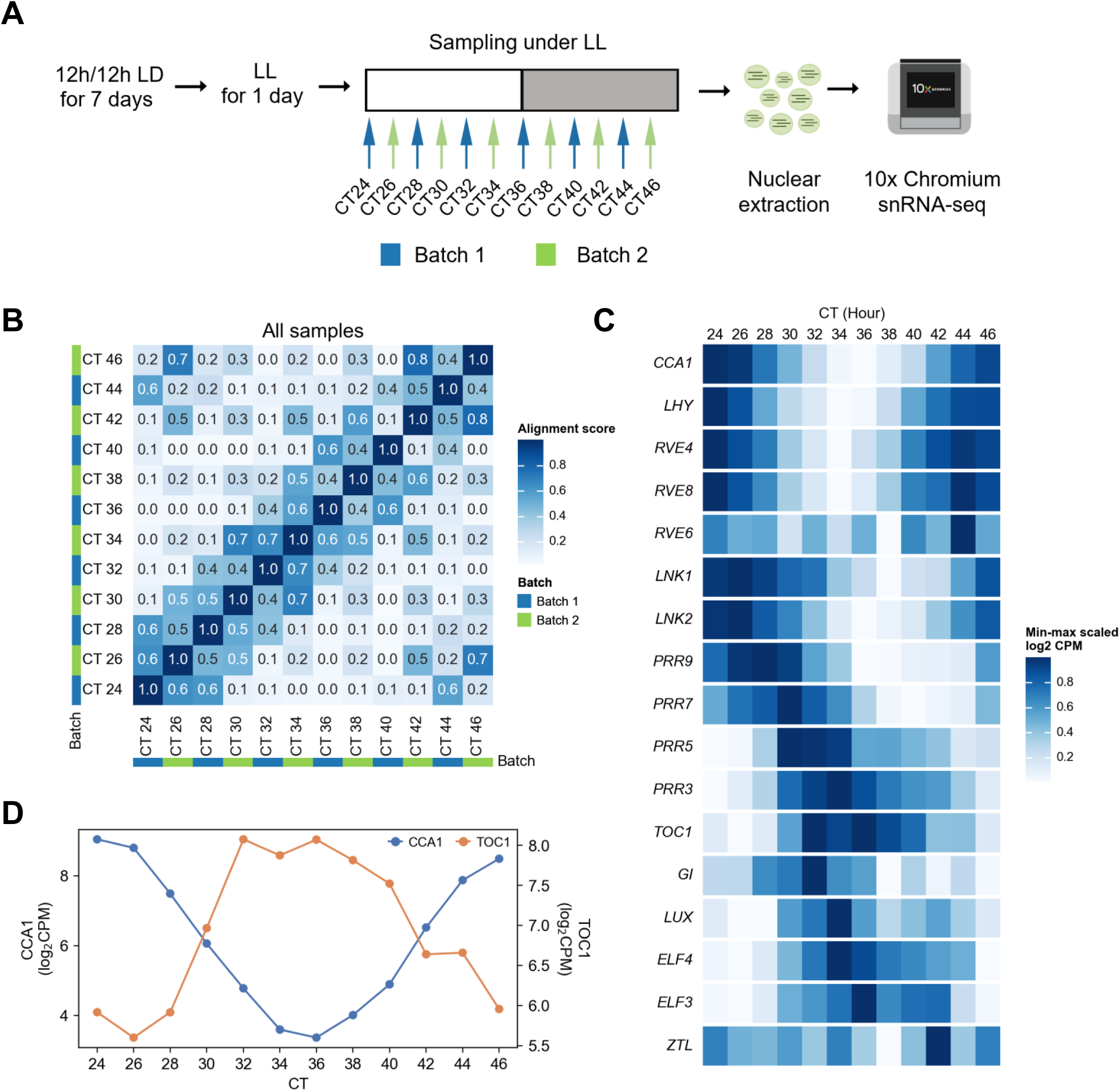
Comprehensive snRNA-seq analysis of transcripts rhythmic accumulations in Arabidopsis seedlings. (A) Experimental design and methodology. Schematic representation of the experimental setup for circadian entrainment, sampling, and snRNA-seq processing. Seedlings were first entrained under a 12-hour light/12-hour dark cycle, followed by a 24-hour shift to continuous light conditions. The snRNA-seq library construction using the 10X Genomics platform was conducted in two separate batches, with subsequent, high-throughput sequencing, and detailed data analysis. (B) Cross-sample correlation analysis. The heatmap presents alignment scores between each sample pair, where these scores represent the proportion of cells identified as mutual nearest neighbors across datasets. Higher scores signify greater transcriptomic similarity between datasets. (C) Heatmap showing the min-max normalized average expression profiles of core clock genes across 12 circadian time points. (D) Line plot illustrating the transcripts accumulation of *CCA1* and *TOC1* across 12 samples, with each point representing the mean expression across all nuclei in a sample.

A recent study in mammal suggests that the circadian clock can affect cell clustering, thus interfering with subsequent analysis^30^. The similarity between samples at adjacent time points suggested the presence of such a phenomenon in our data (Fig. 1B) and the UMAP visualization also indicates that the baseline is different among samples (Supplementary Fig.1). To eliminate this effect and improve clustering results, we evaluated eight data integration strategies, which can be categorized into four groups (Supplementary Fig.2A): (1) Direct application of the Harmony method to integrate data from both batches; (2) Following the approach proposed by Wen et al.^30^, exclusion of clock-regulated genes during the clustering step; (3) Individual application of SCTransform^36^ on each sample to eliminate baseline variations across the data; (4) Employing a procedure analogous to the removal of cell cycle effects^37^, performing SCTransform separately on each batch followed by linear regression to regress out the expression of core clock genes^7^. Next, we evaluated these strategies from the perspective of conserving biological variation and eliminating batch effects. We first performed clustering on each dataset individually (Supplementary Fig.3), and then used the silhouette score to evaluate the performance of these clustering results on the cell distance matrices derived from various strategies (Fig. 2A). This score functioned as an indicator of biological variation conservation. Additionally, we employed the kBET^38^ method to measure the concordance of different samples in the final distance matrix, serving as a metric for evaluating the batch effect removal (Fig. 2A). We observed that the exclusion of clock-regulated genes enhanced the performance of batch effect removal, and removing an equal number of randomly selected genes does not have the impact on batch effect removal (Supplementary Fig.2B). This indicates that circadian rhythms are the main cause of sample heterogeneity. In addition, the removal of clock-regulated genes resulted in a huge decrease in silhouette score, indicating the impaired biological variation (Supplementary Fig.2B). However, the result with the removal of randomly selected genes indicated that this decrease is not solely due to the reduction in the number of genes. It implies the existence of cell-type specific circadian rhythms. Strategy 3 and 4 both exhibited enhancements in both biological conservation and batch effect removal when compared to directly integrating datasets, thus achieving a balance between preserving biological characteristics and removing batch effects. Given the superior performance of strategy 3, we selected it as our final strategy.

**Fig. 2:**
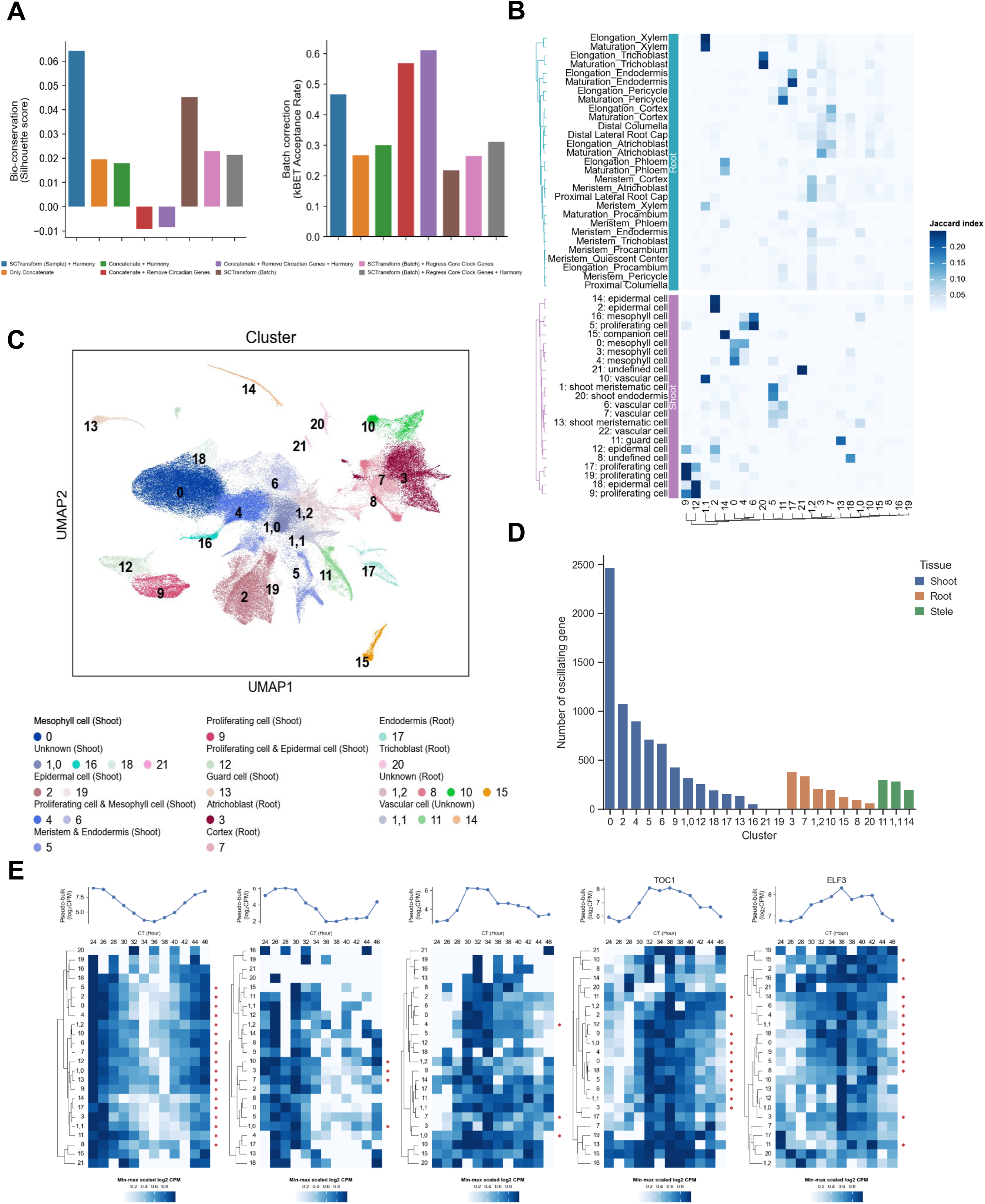
Dissection of cell types and circadian gene expression. (A) Comparison of different integration strategies. We benchmark the different strategies from the perspectives of conserving biological variance (Upper Panel) and removal of batch effects (Lower Panel) (details in the Methods section). For conserving biological variance, we first performed clustering on individual samples and evaluated the clustering results using silhouette score on cell distance matrices obtained from different methods. For mitigating batch effects, we measured the accordance of samples on the nearest neighbor graphs obtained from different methods using the kBET acceptance rate. (B) Heatmap displaying the Jaccard index of cluster-specific genes identified in our snRNA-seq data compared with Arabidopsis shoot and root scRNA-seq datasets. A higher Jaccard index signifies a greater overlap of identified genes between clusters in these datasets. (C) UMAP plot of 22 identified clusters, each labeled according to their origin from shoot, root, or unknown sources (specified in parentheses). (D) Bar plot showing the number of oscillating genes identified within each of the 22 clusters. (E) Top: line plot showing the pseudo-bulk expression levels of a specific gene. Bottom: heatmap representing the mean expression level of the same gene across different clusters at 12 circadian times. An asterisk (*) indicates clusters where the gene is oscillating.

Following data integration and clustering, we identified 22 distinct clusters (Fig. 2C). Each cluster contained nuclei from all time points and both batches with minimal variance (Supplementary Fig. 4). To annotate the identities of each cluster, we first identified cluster-specific expression genes (Supplementary Fig. 5 and Supplementary Table 2), then constructed a pseudo-seedling single-cell dataset by merging two public single-cell root and single-cell shoot datasets^39,40^. Within this synthesized dataset, we identified genes specifically expressed in each cluster and employed the Jaccard Index as a metric to assess the overlap between these specifically expressed genes in our dataset and the pseudo-seedling single-cell dataset. This method guided our annotation, assigning cell types to clusters based on the highest overlap in gene expression (Fig. 2B). Through this process, we were able to identify various cell types from mesophyll, stomata, epidermis, cortex, endodermis, vascular bundles, and the meristem (Fig. 2B). Furthermore, we were able to distinguish the root or shoot origin for all clusters, except for those pertaining to vascular tissue (Fig. 2B), aligning with a previous report that vascular cells from shoots and roots tend to cluster together^39^. Taken together, we effectively reveal cell heterogeneity and classify the major cell types in Arabidopsis seedlings.

### Analyzing clock gene expression in different cell types

To explore circadian rhythm variations across different cell types, we collected 13,256 known clock-regulated genes identified from three global studies^41–43^. Using the JTK-cycle algorithm^44^, we assessed the rhythmicity of these genes across the 12 time points. Our analysis revealed that approximately 40% (5,296/13,256) of clock-regulated genes displayed transcript rhythmic accumulation in at least one cluster (Supplementary Table 3). We found that shoot clusters demonstrated a higher prevalence of oscillating genes compared to root clusters, with mesophyll cells harboring roughly 2,400 oscillating genes (Fig. 2D). Next, we compared the oscillation patterns of core clock genes across different cell types. Our findings revealed that these genes exhibited different oscillation patterns across a variable number of clusters (Supplementary Fig. 6). We selected *CCA1*, *PRR9*, *PRR5*, *TOC1*, and *ELF3* as representative clock genes, which displayed a sequential expression pattern from subjective dawn to midnight (Fig. 2F). Although these genes were detected in nearly all clusters, the extent of their oscillation across clusters showed considerable variation, with the number of clusters displaying rhythmicity spanning from 3 to 18. Specifically, *CCA1* oscillated in 18 clusters, while *PRR9* and *PRR5* were confined to oscillations in only 4 and 3 clusters, respectively (Fig. 2F). This diversity partially underscores the heterogeneity of the circadian clock across cell types. The phase and amplitude analysis of the 24-h rhythms for each cluster revealed that several gene expressions including *CCA1*, *LHY*, *PRR3*, *PRR7*, and *TOC1* showed more robust oscillation in a mesophyll subcluster (cluster 0) compared with other cell types (Supplementary Fig. 7A). In addition, we also observed phase shifts of the circadian rhythms, such as the transcript accumulation of *TOC1*, which display an approximately 2-hour phase delay in shoot epidermal cells relative to other shoot cell types (Supplementary Fig. 7B). Furthermore, we observed considerable variations in phase and amplitude among different vascular cell types for rhythmic expression of clock genes in multiple vascular clusters (Supplementary Fig. 7). Collectively, these findings offer clues for cell-type specific controls within the Arabidopsis clock system.

To corroborate the oscillatory patterns found in our dataset, we generated transgenic lines expressing luciferase (LUC) and β-glucuronidase (GUS) encoding genes driven by the promoters of *RBCS2B* (*AT5G38420*) and *AT1G66100*, respectively. This enabled us to spatially map expression locations and concurrently monitor the rhythmicity of these genes (Fig. 3). We measured the expression levels of these genes under LL conditions and observed a high degree of reproducibility across two biological replicates for each gene (Fig. 3C and 3D). The promoter activity of these genes exhibited a largely similar oscillating pattern with the snRNA-seq data (Fig. 3A-D). It is worth noting that the promoter activity measured at the protein level by LUC assay may not always be the same as RNA-seq measured at the transcript level, and a certain degree of discrepancies are to be expected. In addition, histochemical GUS assays for tissue specificity also aligned with our cell type annotations, supporting the accuracy of our cluster classification (Fig. 3G and 3H). The expression of *RBCS2B*, as indicated by GUS staining, was predominant in shoot-specific cell types (Fig. 3G), corroborating our snRNA-seq findings (Fig. 3A and 3E). Moreover, GUS staining of the cluster-specific gene *AT1G66100* confirmed its epidermal localization, which was in agreement with our initial cluster classification (Fig. 3F and 3H). Collectively, these validations underscore the reliability of snRNA-seq for capturing the intricate spatiotemporal expression dynamics of clock-regulated genes within Arabidopsis.

**Fig. 3:**
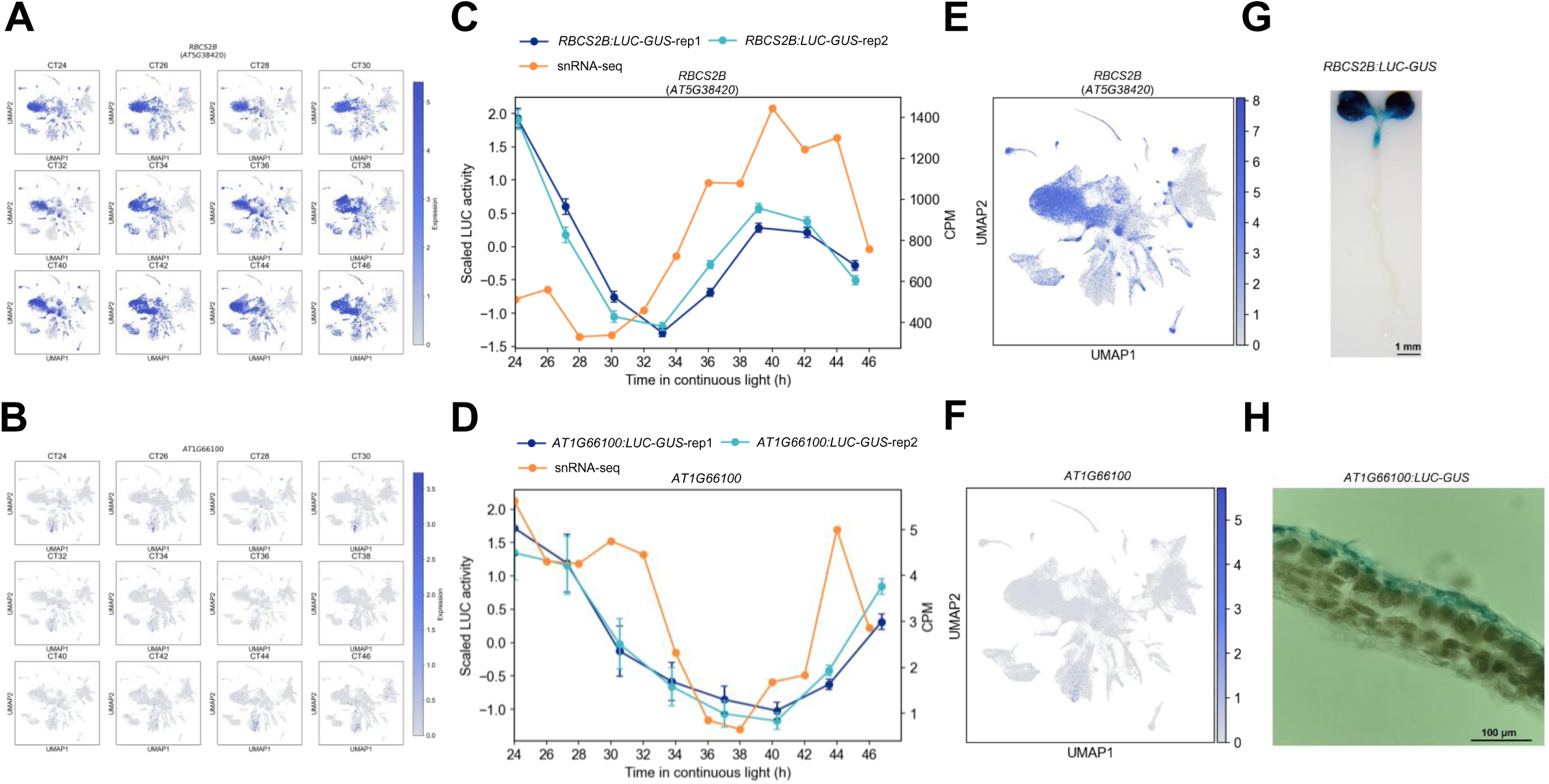
Spatiotemporal validation of oscillatory gene expression patterns. (A), (B) UMAP plots showing the oscillatory expression of *RBCS2B* (*AT5G38420*) (A) and *AT1G66100* (B) across 12 snRNA-seq libraries (C), (D) Line plots showing the average rhythms of *RBCS2B*:*LUC-GUS* luminescence (C), and *AT1G66100*:*LUC-GUS* luminescence (D) in transgenic plants (blue) with two biological replicates. These rhythms are compared with the pseudo-bulk expression from snRNA-seq data (orange), highlighting similarities and discrepancies between in vivo luminescence and sequencing data. (E), (F) UMAP plots representing the specific expression patterns of *RBCS2B* (*AT5G38420*) (E) and *AT1G66100* (F) within the snRNA-seq dataset. (G), (H) Histochemical GUS assays of transgenic seedlings harboring *RBCS2B*:*LUC-GUS* (G) and *AT1G66100*:*LUC-GUS* (H) reporter constructs, demonstrating specific expression patterns.

### Three shoot cell types exhibit synchronized oscillation

In a comparative analysis of gene oscillation across different cell types, we employed the S-MOD method^45^, to statistically assess the overlap of oscillating genes (Fig. 4A). This analysis delineated a clear demarcation between shoot and root cell types, with a pronounced intra-tissue oscillating gene congruence within shoot tissues (Fig. 4B). Furthermore, a subset of shoot cell types – mesophyll, proliferating, meristematic, endodermal cells (clusters 4, 5 and 6) – exhibited substantial relatedness of oscillating gene content (Fig. 4B). Subsequently, pairwise phase analysis of the common oscillating genes identified from these clusters was performed (Fig. 4C). Our findings identified 255, 270, and 208 common oscillating genes across the comparisons of clusters 5 to 4, 6 to 4, and 6 to 5, respectively, with 49, 19, and 23 of these genes exhibiting substantial phase shifts. Predominantly, most of the genes with phase differences showed significant phase lag mainly in cluster 5 compared with cluster 4 and cluster 6, suggesting a delayed oscillatory phase in endodermis and meristem cells compared to proliferating and mesophyll cells (Fig. 4C). Conclusively, our study delineates a clear distinction in oscillating gene content between shoot and root cell types, with a subset of shoot cell types exhibiting markedly enhanced oscillator coherence, underscoring a complex, tissue-specific regulation of gene oscillation.

**Fig. 4:**
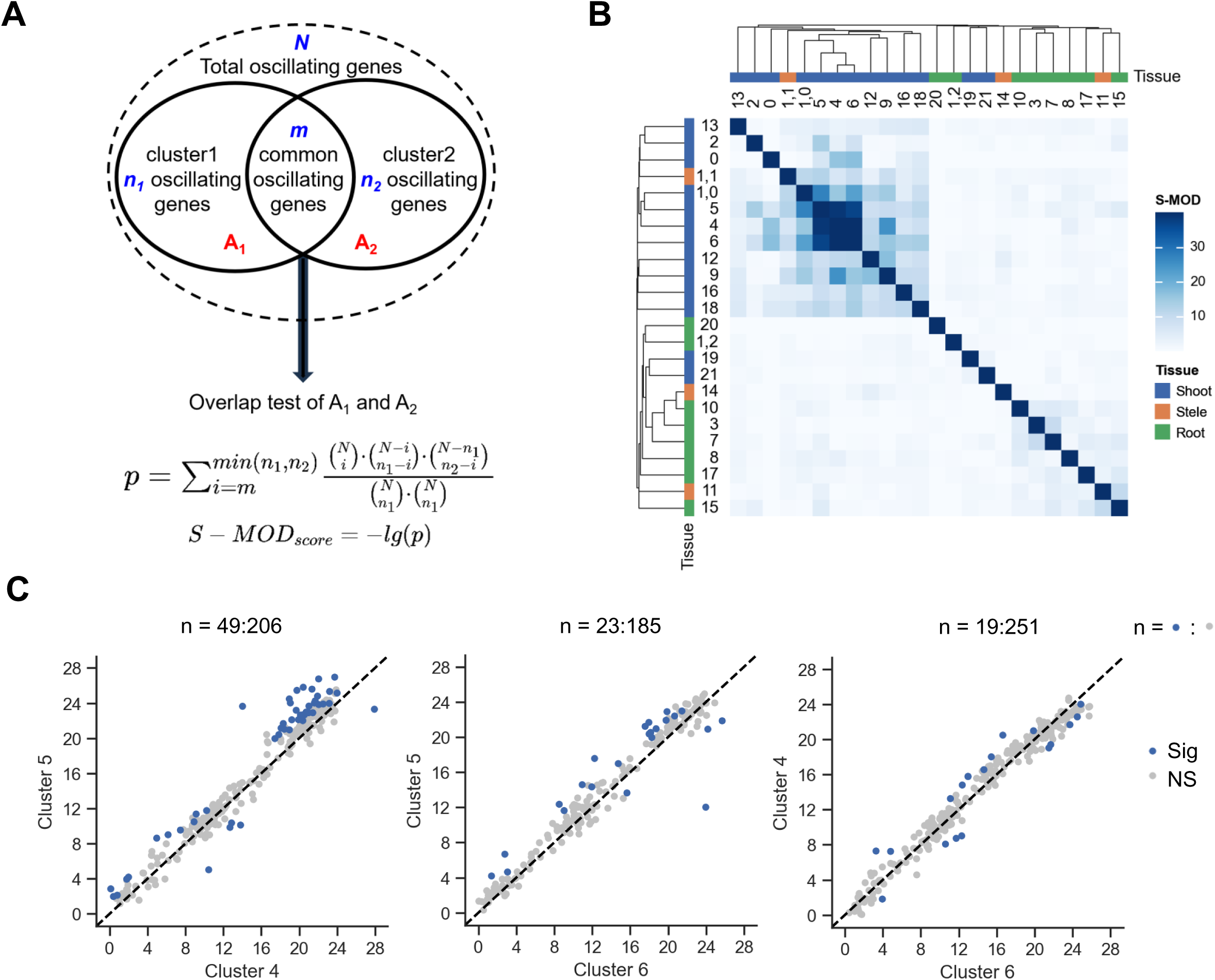
Shoot clusters exhibit significant similarity in oscillating gene contents. (A) Summary of the S-MOD calculation for pairwise statistical overlap between oscillating genes. The significance of the overlap (p-value) between two clusters was calculated considering various parameters: n1 and n2 represent the number of oscillating genes in the two respective gene sets (A1 and A2), m is the count of common oscillating genes shared by the clusters, and N denotes the total number of oscillating genes identified in the entire dataset. The S-MOD score is defined as the negative logarithm of the p-value, offering a clear measure of the statistical significance of the overlap. (B) Heatmap displaying the S-MOD scores for pairwise comparisons of oscillating genes across 22 clusters. To facilitate easy differentiation, clusters originating from shoot, root, and stele are color-coded in blue, orange, and green, respectively. (C) Pairwise phase analysis between cluster 4, 5 and 6. The number of genes (n) showing significant phase shifts is indicated above each plot, with the blue dots representing genes with statistically significant phase differences (Sig), and grey dots indicating non-significant phase relationships (NS).

### snRNA-seq identifies the circadian clock-controlled genes in various cell types

Observing the rhythmic expression of multiple core clock genes across various cell types, we next quantified the number of clusters where each gene oscillated. Our findings indicated that approximately 61% (3,238/5,296) of genes only oscillated in a specific cluster (Fig. 5A), ∼ 10% (341/3,238) of which are cluster-specific genes (Supplementary Fig. 8). This finding of cell-type specific oscillation is consistent with a previous bulk microarray study in Arabidopsis, which indicated limited clock-regulated gene overlap between mesophyll cells and the vasculature ones^19^. A similar phenotype has also been reported in the Drosophila clock system, where the oscillating genes exhibit minimal consistency across cell types^31^.

**Fig. 5:**
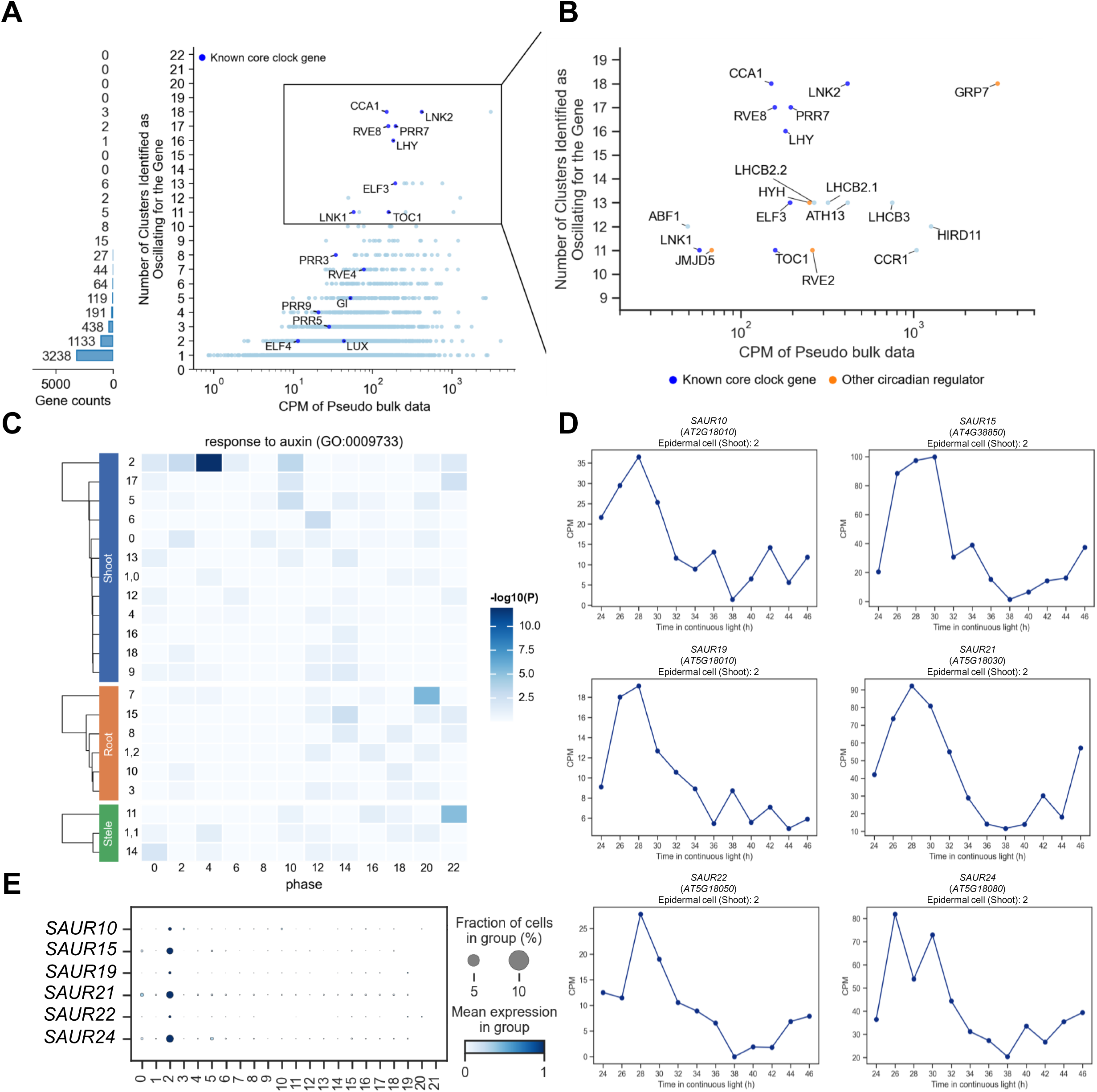
Identification of cell-type specific oscillating genes. (A) Left: Bar plot illustrating the quantity of oscillating genes identified across varying numbers of clusters. Right: Scatter plot representing the correlation between the pseudo-bulk expression level of each oscillating gene and the corresponding count of clusters where the gene is identified as oscillating. Core clock genes are highlighted in blue. (B) An enlarged section of Fig.5A, which allowing for a more detailed analysis of genes that oscillate in more than 10 clusters. Known core clock genes are highlighted in blue; genes regulating clock gene expression colored in orange. (C) Heatmap displaying the enrichment of the Gene Ontology (GO) term ’response to auxin’ (GO:0009733) across different cell types. For detailed statistical data and analysis, please refer to Supplementary Table 4. (D) Line plots indicating the expression of six *SAUR* genes in shoot epidermal cells (cluster 2) in snRNA-seq data. (E) Dot plot representing the expression profiles of six *SAUR* genes across various clusters. The size of each dot correlates with the percentage of cells expressing the gene within a cluster, while the color intensity reflects the gene expression level.

Further Gene Ontology (GO) analysis of oscillating genes peaking at different phases within each cluster uncovered both common and distinct functions among different cell types (Supplementary Table 4). Specifically, genes involved in light response and photosynthesis were enriched among the shoot cell types (Supplementary Fig. 9). Moreover, our analysis revealed unique functional terms that were enriched in specific cell types. For instance, a GO term related to auxin response was predominantly enriched in shoot epidermal cells around CT4 (Fig. 5C), involving a set of genes associated with the auxin signaling pathway, including six *Small Auxin-Upregulated RNA* (*SAUR*) family genes (*SAUR10*, *15*, *19*, *21*, *22*, *25)*. These genes were found to oscillate only in shoot epidermal cells, peaking in the early morning hours (Fig. 5D), and exhibited cell-type specific expression patterns in these cells (Fig. 5E). These findings underscore the strength of our data in identifying cell-type specific clock-regulated genes.

### A data-driven approach for identifying key circadian regulators

Our findings further indicate that core clock genes, including *CCA1* (18), *LNK2* (18), *PRR7* (18), *LHY* (16), and *RVE8* (17) are oscillating in most of the 22 clusters (Fig. 5A). Then, shifting focus to the 19 genes that oscillated in over 10 clusters, we found that half of them are reported as circadian regulators, including 8 core clock genes, and other 4 genes including *glycine-rich RNA-binding proteins 7* (*GRP7*)^46^, *jumonji domain containing 5* (*JMJD5*)^47^, *RVE2*^48^, and *HY5-HOMOLOG (HYH)*^49^ (Fig. 5B). Hence, genes found to oscillate in multiple cell types are indeed genes that play a critical role in the regulation of circadian rhythms. Considering that clock components exhibit significant evolutionary conservation across angiosperms^50,51^, our snRNA-seq data on circadian expression can be leveraged beyond Arabidopsis research and serve as a robust tool to pinpoint circadian regulators in a diverse range of plant species. Moreover, for the many plant species in which the circadian components remain partially unclear^52–54^, our time-series snRNA-seq approach could be applied to quickly generate a potential gene list of key circadian regulators ranked by the number of cell types that they are oscillating in.

Augmenting the utility of our study, we have curated an online website that provides easy access to gene expression data from our research (Fig. 6). The portal facilitates searches using gene IDs, with immediate visualization and access to the expression profile of each gene, including UMAP plots, bar plots, and line plots detailing expression patterns across circadian time points at both the whole plant and cell-type levels (Fig. 6D-G), thus providing a valuable resource for the community and fostering a collaborative environment for furthering circadian rhythm research.

**Fig. 6:**
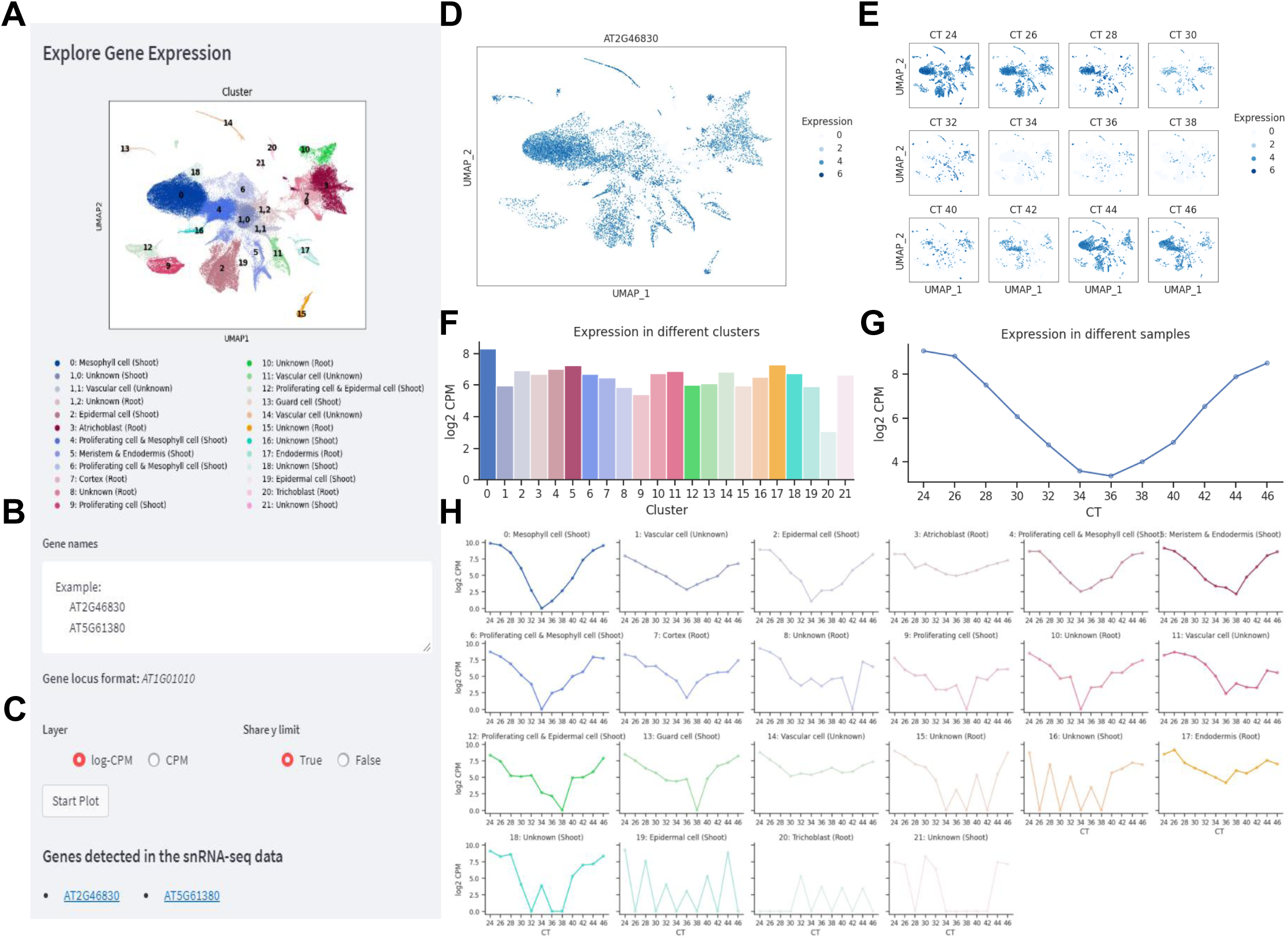
Interactive web portal for exploring the snRNA-seq dataset. (A) UMAP plot representing a visualization of the 22 identified cell types. (B) The interface provides a user-input field for querying specific gene IDs of interest. (C) Users can select the preferred gene expression unit for visualization through the ’Layer’ option, which includes Counts Per Million (CPM) or logarithm base 2 of CPM (log_2_CPM). Additionally, the ’Share y Limit’ feature allows for consistent y-axis scaling across multiple plots, enhancing comparability, as exemplified in the bar plots of Fig. 6F. Genes detected in the snRNA-seq data are shown below. (D) UMAP plot representing the gene expression profiles across 22 clusters. (E) UMAP plots representing the expression profiles across 12 samples. (F) Bar plots representing gene expression profiles within each cluster. (G) Line plot showing the overall expression profiles at the whole seedling level. (H) Line plots showing expression profiles across the clusters.

Our time-series high-throughput snRNA-seq data has provided a new perspective on circadian gene expression within individual cell types of plants. This method represents a significant step forward in understanding the cell-autonomous circadian clock, offering insights into the complexities of circadian regulation. The potential future application of this technique to other plant species could contribute to a broader understanding of circadian rhythms in plant biology and potentially inform agricultural practices.

## Methods

### Plant materials

Plant growth and material harvesting were performed as previously described with minor modifications^55^. The *Arabidopsis thaliana* used in this study was the Columbia-0 ecotype. Seeds were sown on MS agar plates containing 3% sucrose and stratified at 4 ℃ for 2 days, then placed in growth chambers at 22 ℃. Plants were entrained by 12 h light and 12 h dark (12L/12D, 100 μmol m^-2^ s^-1^) photoperiod for seven days and then released into constant light (100 μmol m^-2^ s^-1^) conditions, harvested every 2 hours over a 24-hour period starting at subjective dawn on day 9.

### Tissue-specific LUC-GUS transgenic plant construction, GUS staining, monitoring circadian rhythms

The promoter regions of *RBCS2B* (*AT5G38420*) and *AT1G66100* which encode a PR (pathogenesis-related) protein were amplified using the Phusion High-Fidelity DNA Polymerase (New England Biolabs) from Col-0 genomic DNA with two primer pairs RBCS2B-BglII-PF (5′-CCAGATCTGGAATATTCAATGTTGACTATC-3′) and RBCS2B-Eco91I-PR (5′-CGTCGGTGACCATGGCTTCTTCTTGTTGTTTCTCTTC-3′), AT1G66100-BamHI-PF (5′-GTGGATCCAGTCCATTAATGTCATAAATCTG-3′) and AT1G66100-Eco91I-PR (5′-CGTCGGTGACCATCTTTTGATTGATTAGTTG-3′). PCR products of *RBCS2B* promoter region (2,364 bps) and PCR products of *AT1G66100* promoter region (2,600 bps) were double digested with BamHI and Eco91I or BglII and Eco91I, then the promoter-driven LUC-GusP fusion reporter plasmids were made through insertion of the digested PCR-amplified promoter regions into a modified *pENTR 2B* vector with *LUC-GusP* gene at the multiple cloning site (MCS). Finally, The *promoter:LUC-GusP* constructs were recombined into a modified pH2GW7Δ^56^ from which the 35S promoter had been deleted via Gateway cloning technology. *RBCS2B:LUC-GusP* and *AT1G66100:LUC-GusP* transgenic (T3) lines were used to determine the tissue-specific expression pattern via histochemical GUS reporter activity^57^. For firefly LUC measurement, two reporter transgenic lines were entrained for 10 days in LD (12h light/12h dark, 22°C) before release to the free-running conditions for luciferase activity analysis with an EM-CCD cooled camera system (Andor Technology).

### Nuclei isolation and 10X single-nucleus RNA-seq library preparation

Frozen samples from 9-day-old Arabidopsis seedlings were used for single nucleus preparation. The nuclei were isolated as previously described^58^. In short, the seedlings were chopped in ice-cold 1 x Nuclei Isolation Buffer (NIB, MilliporeSigma, cat. no. CELLYTPN1) supplemented with 1mM dithiothreitol (DTT, Thermo Fisher, R0861), 1 x protease inhibitor (Sigma, 4693132001) and 0.4 U/μl Murine RNase inhibitor (Vazyme, R301-03). Thereafter, the lysate was filtered through a 40 μm strainer and centrifuged at 4 ℃ for 5 minutes at 1,000 g. The supernatant was then removed, and the nuclei pellet was resuspended with 500 μl NIB. To further remove the debris, the nuclei suspension was filtered with a 20 μm strainer and stained with 4,6-Diamidino-2-phenylindole (DAPI, 1 mg/ml) before fluorescent activated cell sorting (FACS). The isolated nuclei were run on an MA900 flow cytometer (SONY) with a 100 μm nozzle and sorted into a tube containing 1 ml of collection buffer (1× PBS supplemented with 1% BSA and 0.4 U/μl Murine RNase inhibitor). At least 100,000 nuclei were sorted per sample based on the DAPI signal intensity and the nuclear size. The single-nucleus suspensions were pelleted at 4 ℃ for 5 min at 1,000 g and then resuspended in 45 μl collection buffer. The quality of the nuclei was checked under the microscope. Approximately 20,000 intact nuclei were loaded onto the Chromium Next GEM Chip G, followed by the library construction for Illumina sequencing performed as previously described^59^.

### Single-nucleus RNA-seq data analysis

We first utilized Cell Ranger for preprocessing the raw Fastq files. All parameters are set to their default values, except for the “--include-introns” flag to accommodate the high intron retention ratio feature of the nucleus transcriptome. Next, we use scdblfinder^60^ to eliminate potential doublets and SoupX^61^ to remove ambient RNA contamination, thereby obtaining an initially filtered matrix. The obtained matrix was then subjected to additional filtering using the median absolute deviation (MAD) method recommended in the Single-Cell Best Practices^62^. In this step, cells with gene counts, UMI counts, and organelle transcript proportions exceeding 5 MAD were classified as outliers. Furthermore, cells with an organelle transcript proportion exceeding 10% are further removed from the matrix.

After completing quality control steps for each library, we merge the filtered results to obtain the final gene expression matrix. Subsequently, we tested multiple integration strategies to identify a suitable method for our data: (1). Performing PCA directly on the concatenated dataset, either with or without the removal of known circadian genes^41–43^. This is followed by the construction of a nearest neighbor graph and clustering. (2). Applying Harmony^63^ to the concatenated dataset to alleviate batch effects. This is followed by the construction of a nearest neighbor graph, either with or without the removal of known clock-regulated genes. (3). Performing SCTransform separately on each batch, then removing or retaining the expression of core clock genes (achieved by specifying the “vars.to.regress” parameter). Next, we either perform Harmony or refrain from it on the resulting Pearson residuals. (4). Performing SCTransform separately on each sample to remove the baseline difference, followed by the Harmony method.

Among all the methods, the number of highly variable genes was set to 3,000. PCA and the construction of a nearest neighbor graph were performed using the functions provided in Scanpy^64^ with default parameters. In addition, we also performed clustering on each sample separately. Apart from setting the number of highly variable genes to 3,000, all other parameters were consistent with the tutorial of Scanpy. The clustering results will be evaluated using the silhouette coefficient, implemented with the functions wrapped in sklearn^65^, on the cell distance matrix obtained from the aforementioned methods. We also evaluated the effectiveness of batch effect removal using the kBET^38^ method provided in scIB^66^.

After the clustering method is determined, we employ the UMAP method^67^ (scanpy.tl.umap,zmin_dist=0.3) to obtain a two-dimensional embedding of the cells, and the Leiden method^68^ (scanpy.tl.leiden, resolution=0.4) to obtain clustering results. Upon careful examination of the expression patterns of known cell type marker genes, we observed that Cluster 1 encompassed cells originating from various cell types. Consequently, we proceed to perform further clustering within this cluster (scanpy.tl.leiden, restrict_to=Cluster 1, resolution=0.1). Subsequently, we utilized the cellex method to identify cluster-specific genes^69^. This method integrates various indicators such as gene expression ratios, T-test results, and gene enrichment scores to calculate a specificity score for each gene in different clusters, ranging from 0 to 1 and we use a threshold of 0.9 as the final cutoff. In order to annotate the identity of each cluster, we combined the Root data from Shahan et al. with the Shoot data from Zhang et al^39,40^. Subsequently, we also utilized the cellex method to identify cluster-specific genes in this pseudo-seedling dataset. To assess the similarity between our clusters and the clusters in the pseudo-seedling dataset, we employed the Jaccard Index as a measure.

To determine which genes among the reported clock-regulated genes exhibit oscillation in our dataset, we collected clock-regulated genes identified in three different sources of transcriptomic data. We then followed the approach proposed by Wen et al. to identify the subset of genes oscillating in our data. Initially, we merged all cells within each cluster to obtain pseudo-bulk data. Next, after performing CPM normalization and log2- transformation, we used the JTKcycle method wrapped in the metacycle package to calculate the corresponding p-values^44^. In this step, the circadian periods were set to 24 hours. Genes with adjusted p-values below 0.05 were considered as oscillating. We then employed a nonlinear regression to fit the cosine curve and determine the phase, amplitude, and corresponding standard deviation errors of these oscillating genes. We also utilized the z-score to compute the statistical significance of phase and amplitude as described by Yan et al^30^. In general, the z-score is obtained using the formula: 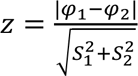, where φ represents phase or amplitude, and S represents standard deviation. Subsequently, the two-tailed p-value is calculated using the scipy.stats.norm.cdf function.

## Data Availability

The raw sequencing data generated in this study were deposited in the China National Center for Bioinformation with accession PRJCA021408 (https://ngdc.cncb.ac.cn/gsa/s/eEfiBu1e for reviewer link).

## Code Availability

The source code to reproduce this project can be accessed at https://github.com/ZhaiLab-SUSTech/circadian_notebooks.

## Acknowledgments

The group of J.Z. is supported by the National Natural Science Foundation of China (32325031), National Key R&D Program of China Grant (2019YFA0903903); Y.L. is supported by the National Natural Science Foundation of China (32100444) and Natural Science Foundation of Guangdong Province of China (2023A1515011997); the Shenzhen Fundamental Research Program (JCYJ20210324105202007); and the National Natural Science Foundation of China to Q.X. (32170259, 31670285) and X.X. (U1904202, 32170275).

## Author Contributions

Y.Q., J.Z., X.X., and Y.L. designed the experiments. Y.Q., Y.L., X.Z and S.G performed the experiments. Y.Q., Z.L., X.Z and Y.L. analyzed the data. Y.G generated the constructs. Y.Q., Z.L., J.Z. and X.X. wrote the manuscript. J.Z., M.A.N. and X.X. oversaw the study. B.L. and Q.X. provided conceptual insight.

