## Supplementary Figure for "*A Day in the Life of Arabidopsis:* 24-Hour Time-lapse Single-nucleus Transcriptomics Reveal Cell-type specific Circadian Rhythms"

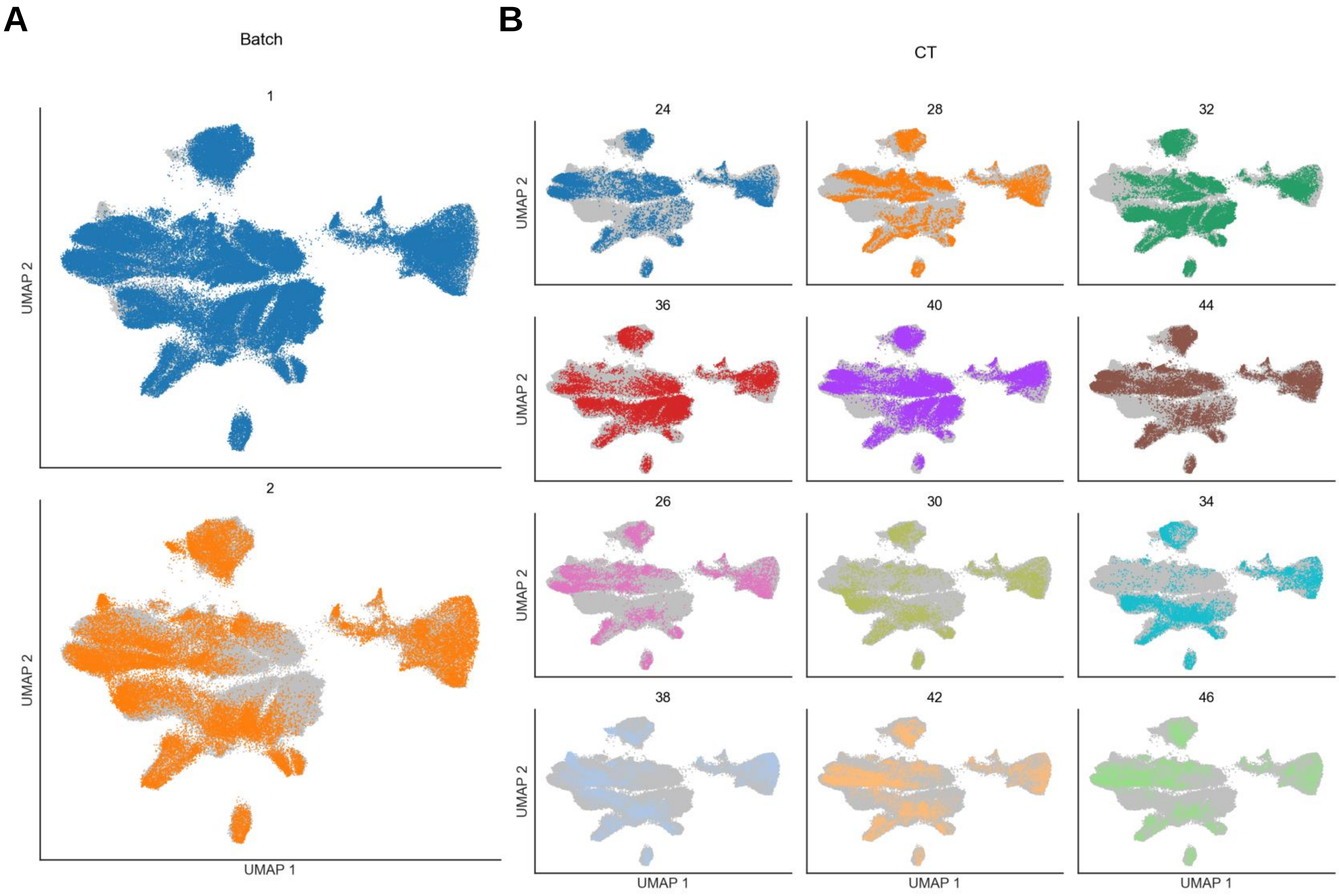


**Supplementary Fig. 1: The UMAP visualization of concatenated snRNA-seq data.**

(A) UMAP plots delineating nuclei from different batches. Nuclei from batch 1 are represented in blue, while those from batch 2 are in orange.

(B) UMAP plots displaying nuclei from 12 different samples, annotated with their respective circadian times (CT).

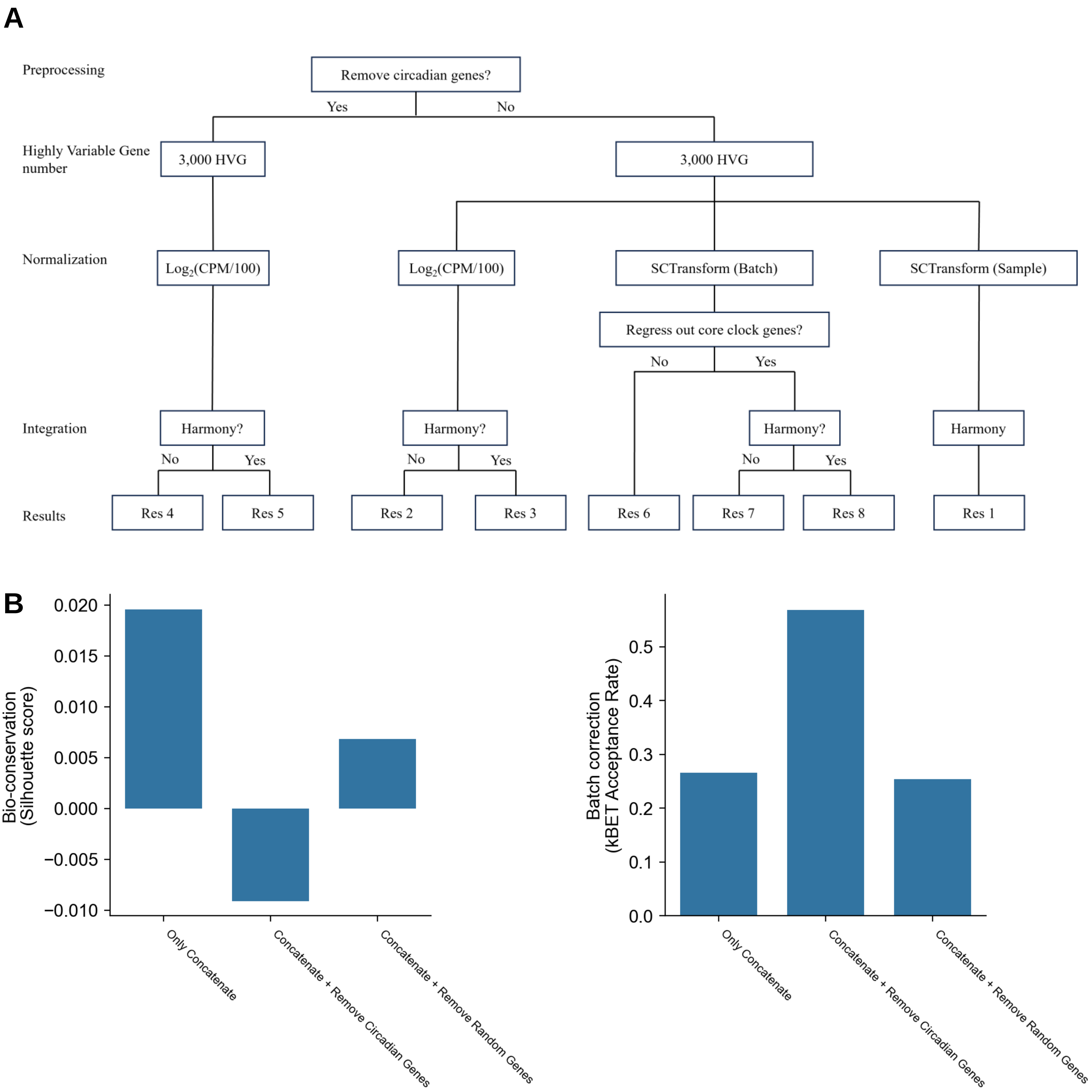


**Supplementary Fig. 2. Benchmarking different data integration strategies.**
(A) Flow charts of eight different strategies for data integration.
(B) Comparison between the removal of clock-regulated genes and the random removal of the same number of genes.

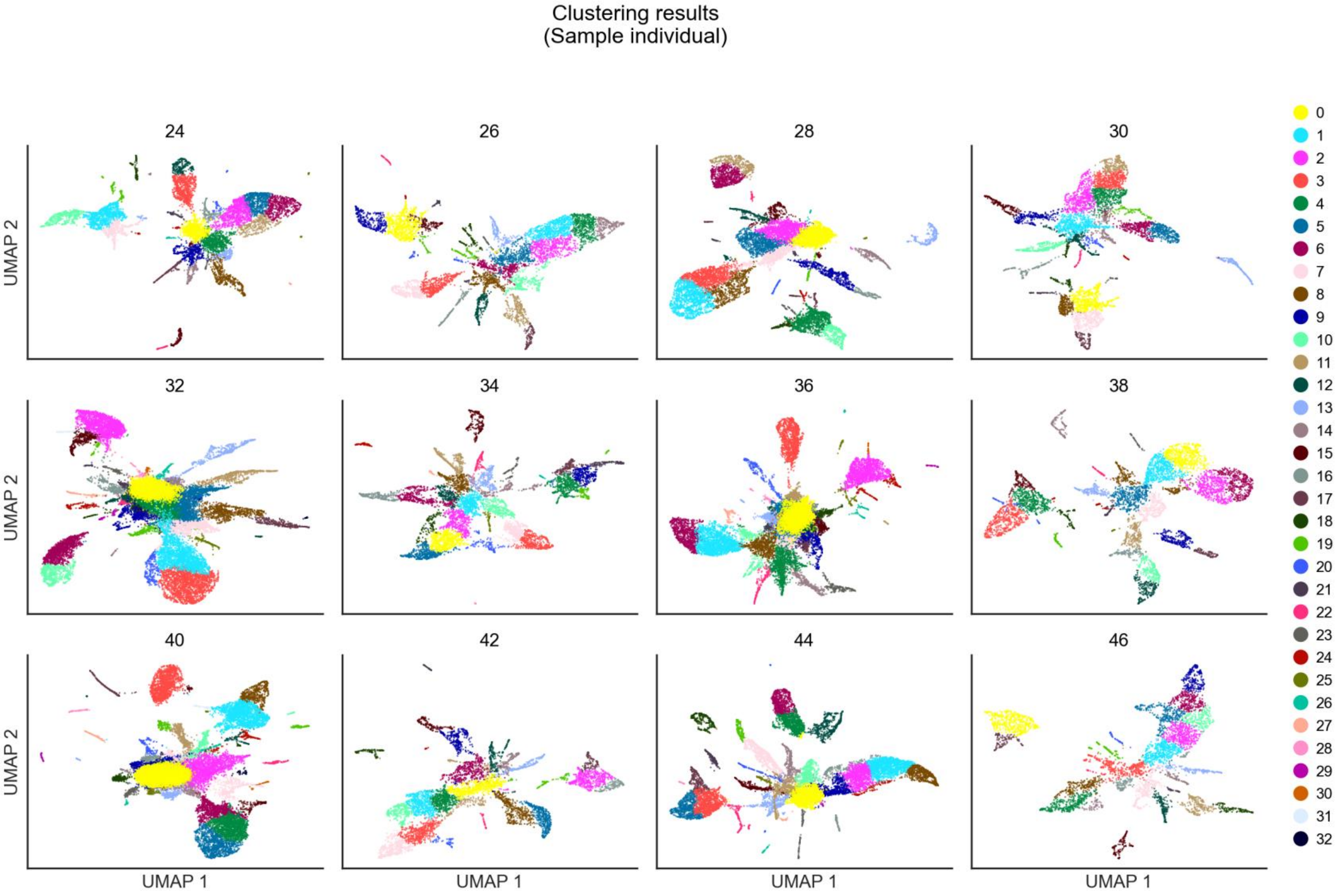


**Supplementary Fig. 3: Clustering and UMAP visualization were conducted separately for each dataset.** The parameters used in this process were kept consistent for all samples (3,000 highly variable genes and a clustering resolution of 1.0). However, due to the independent analysis of the samples, there is no assurance that cells with the same cluster label in different samples correspond to the same cell type.

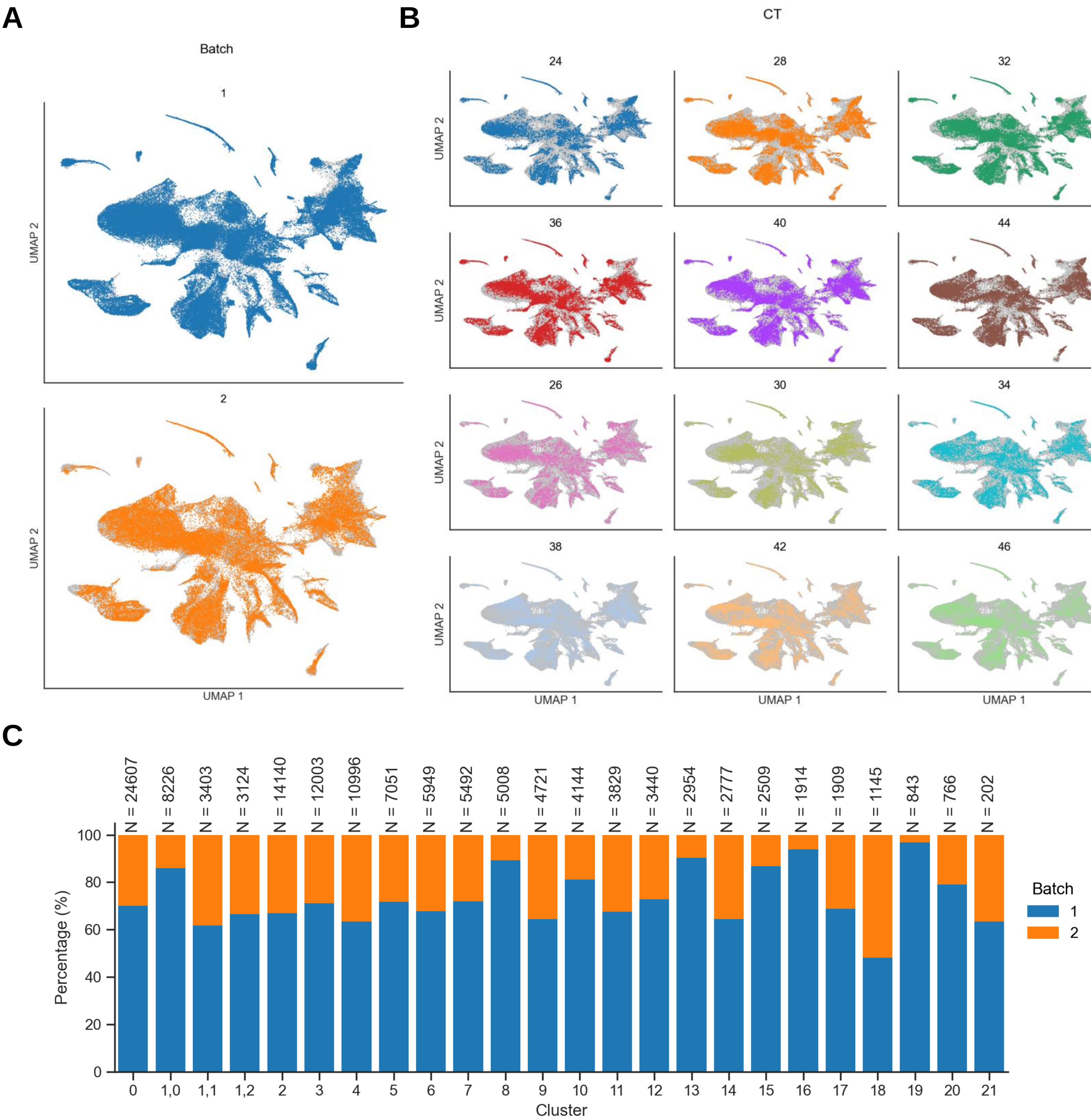


**Supplementary Fig. 4: Integration of the snRNA-seq data.**
(A) UMAP plots of nuclei from batch 1 (blue) and batch 2 (orange) after data integration.
(B) UMAP plots of nuclei from 12 samples after data integration.
(C) Bar plots showing the proportion of nuclei from two batches in each cell type.

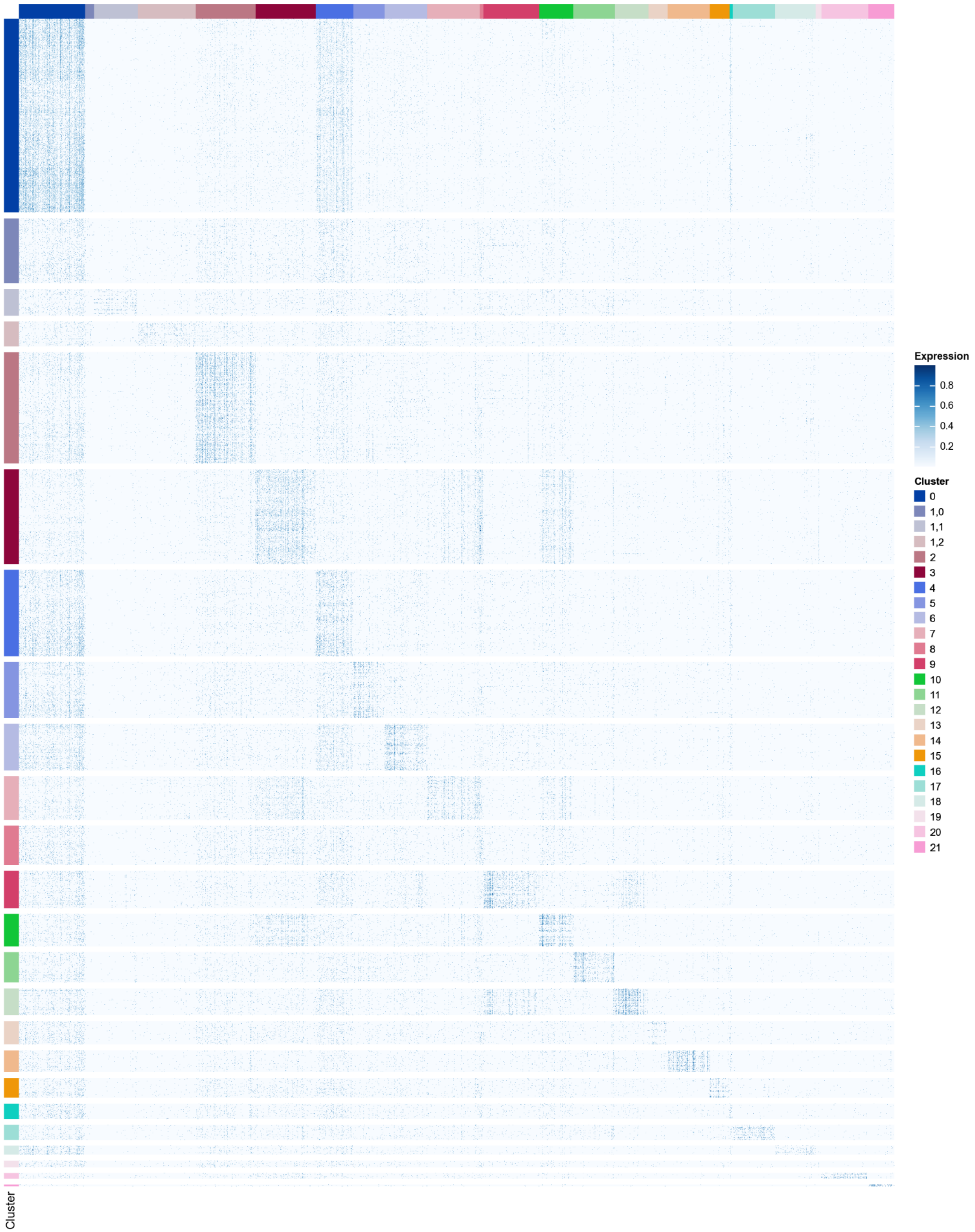


**Supplementary Fig. 5: Expression of cluster-specific genes**. Heatmap showing the expression pattern of cluster-specific genes across cell types.

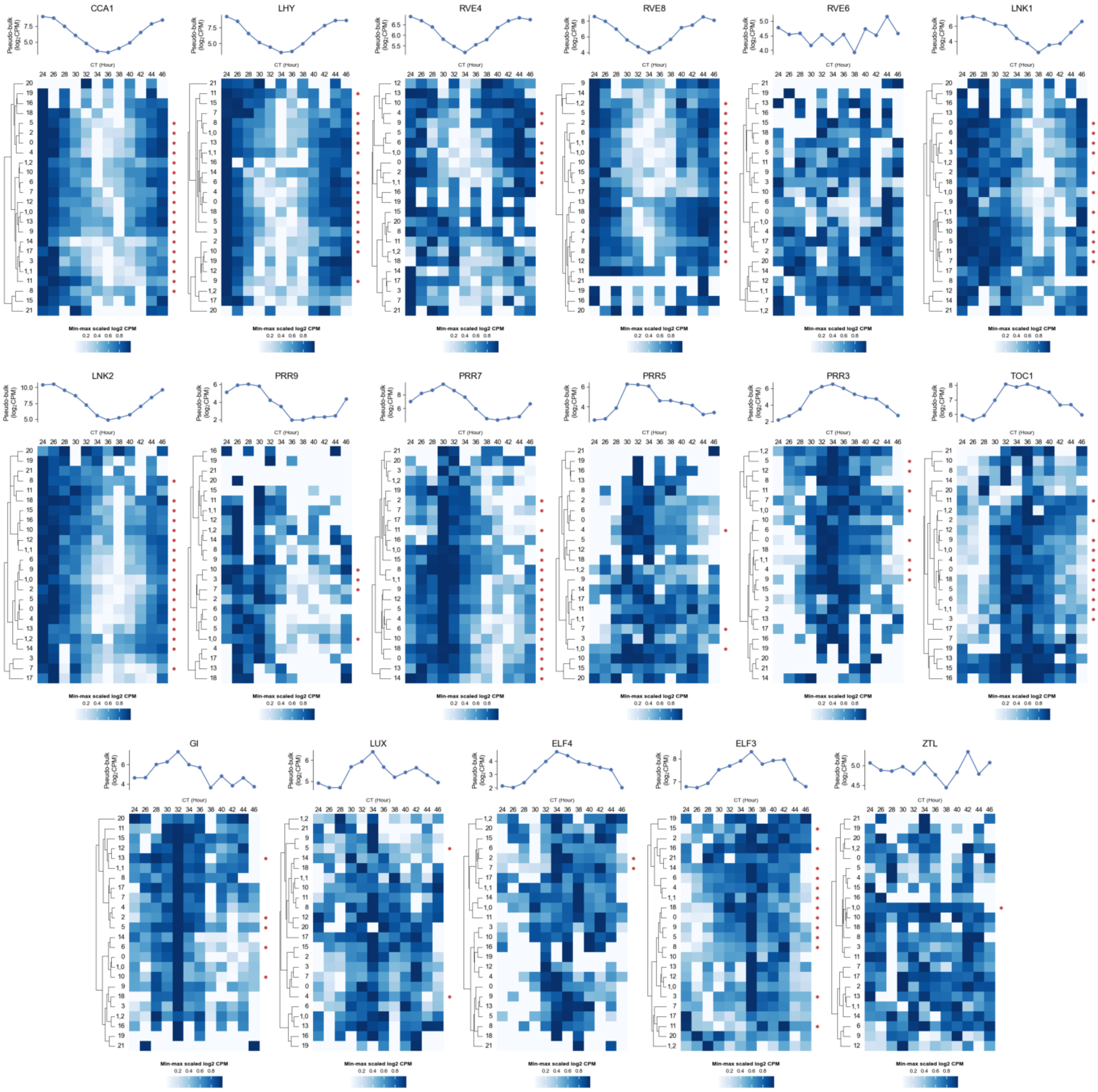


**Supplementary Fig. 6: Core clock gene expression across cell types.** Upper Panel: line plots delineating pseudo-bulk gene expression profiles across 12 samples. Lower Panel: heatmap plots showing the expression patterns of these genes across various cell types. An asterisk (*) indicates clusters where the gene is oscillating.

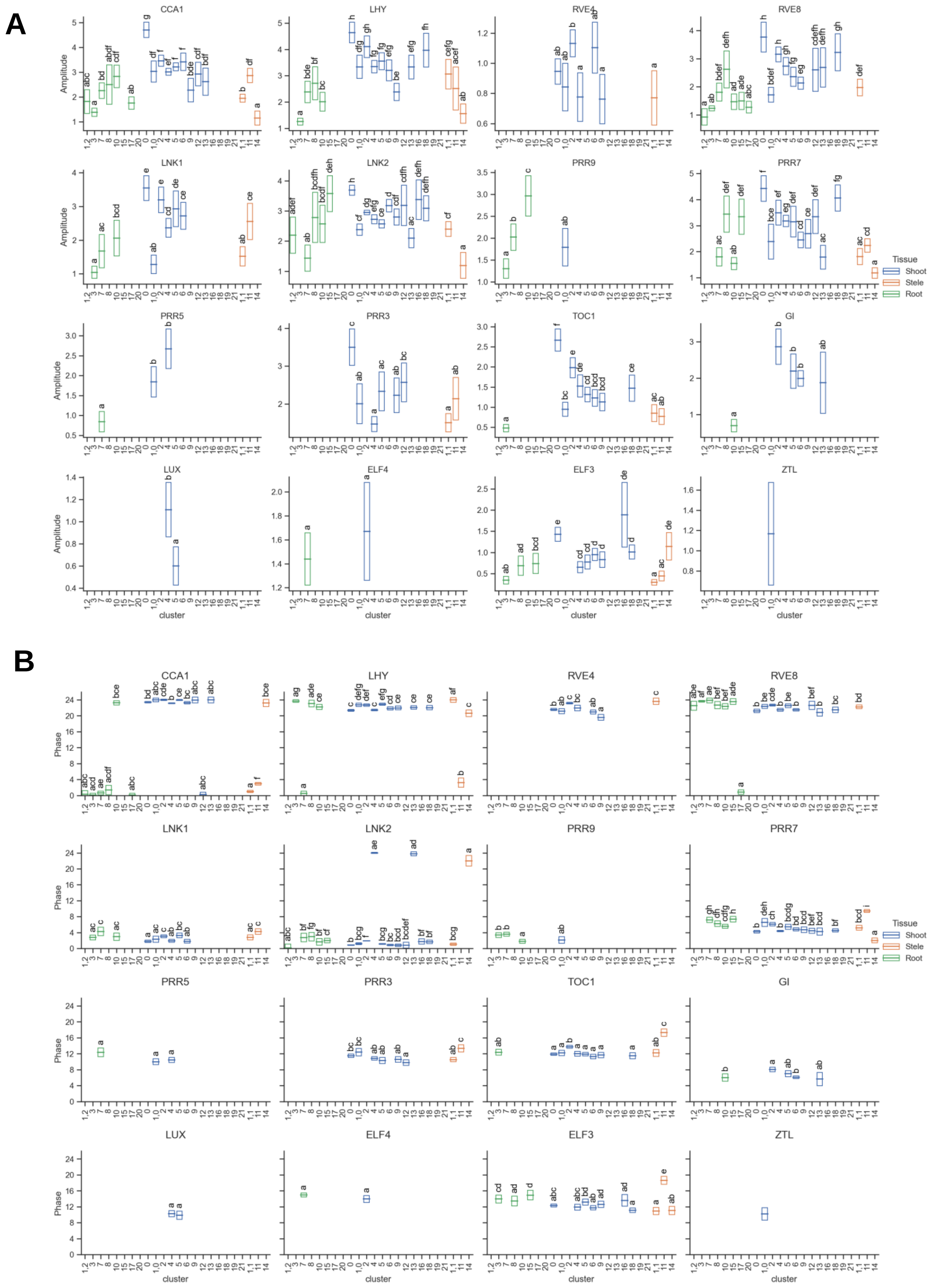


**Supplementary Fig. 7: Phases and amplitudes of core clock genes across cell types.** The rectangle illustrates the estimated amplitude (A) and phase (B) range, denoted by $\varphi \pm$s.e.m.. P value was calculated by z-score based methods (See methods). Different letters above the rectangle indicate statistically significant differences in amplitude (A) and phase (B) ($p < 0.05$), while identical letters signifies no significant differences detected.

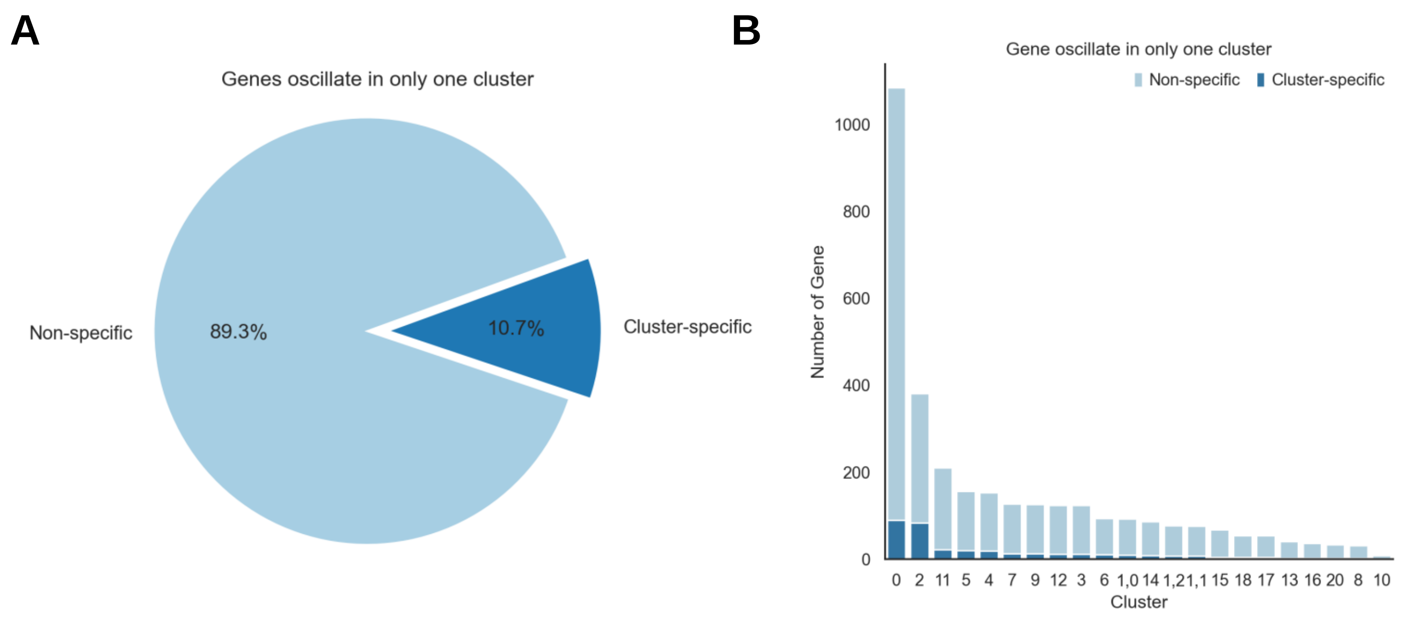


**Supplementary Fig. 8: Composition of genes oscillate in only one cluster.**

(A) The pie chart quantifies the oscillating genes that are exclusive to a single cluster, with 10.7% displaying cluster-specific expression and 89.3% showing non-specific expression patterns.

(B) Bar chart details the distribution of genes oscillate in only one cluster across each individual cluster.

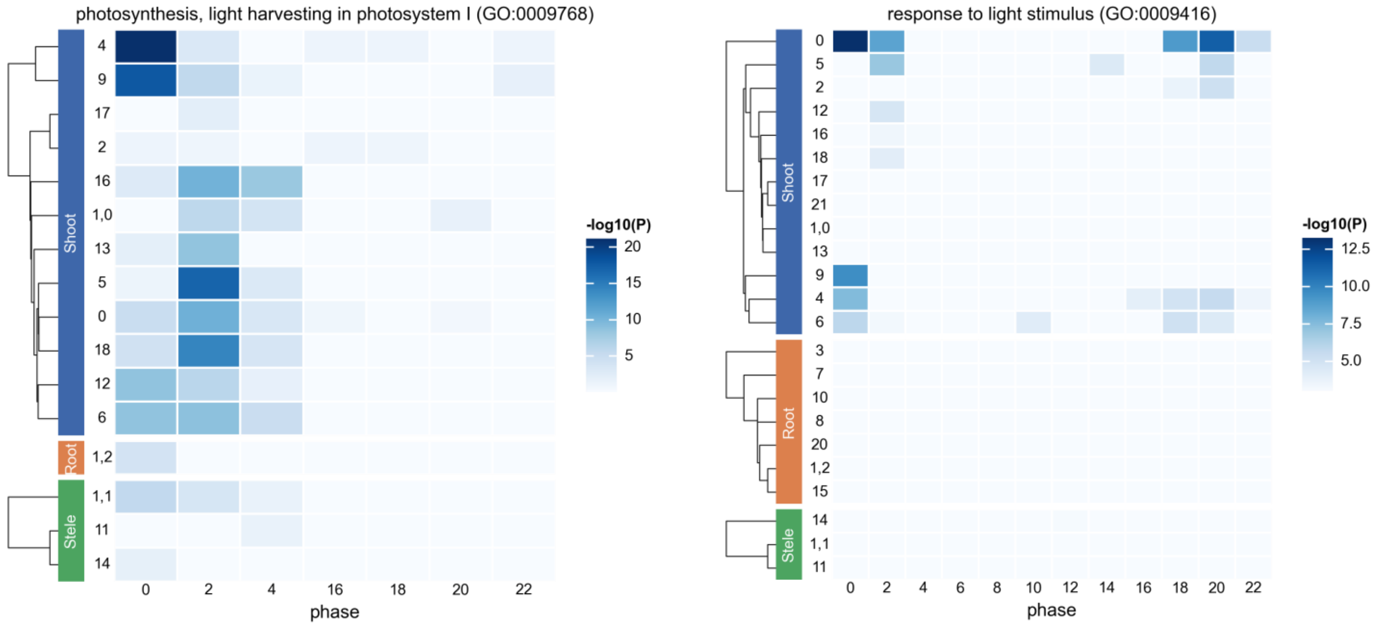


**Supplementary Fig. 9: GO analysis of oscillating genes in different cell types.** Heatmap displaying the enrichment of the GO term ‘photosynthesis, light harvesting in photosystem I’ (GO:0009768) and ‘response to light stimulus’ (GO:0009416) across different cell types. For detailed statistical data and analysis, please refer to Supplementary Table 4.
